# Impact of low-frequency coding variants on human facial shape

**DOI:** 10.1101/2020.07.30.227769

**Authors:** Dongjing Liu, Nora Alhazmi, Harold Matthews, Myoung Keun Lee, Jiarui Li, Jacqueline T. Hecht, George L. Wehby, Lina M. Moreno, Carrie L. Heike, Jasmien Roosenboom, Eleanor Feingold, Mary L. Marazita, Peter Claes, Eric C. Liao, Seth M. Weinberg, John R. Shaffer

## Abstract

The contribution of low-frequency variants to the genetic architecture of normal-range facial traits is unknown. We studied the influence of low-frequency coding variants (MAF < 1%) on multi-dimensional facial shape phenotypes in 2329 healthy Europeans. We used MultiSKAT o scan the exome for face-associated low-frequency variants in a gene-based manner. Seven genes (*AR, CARS2, FTSJ1, HFE, LTB4R, TELO2, NECTIN1*) were significantly associated with shape variation of the cheek, chin, nose and mouth areas. These genes displayed a wide range of phenotypic effects, with some impacting the full face and others affecting localized regions. The missense variant rs142863092 in *NECTIN1* had a significant effect on chin morphology, and was predicted bioinformatically to be deleterious. *NECTIN1* is an established craniofacial gene that underlies a human syndrome that includes a mandibular phenotype. We further showed that *nectin1a* mutations can affect zebrafish craniofacial development, with the size and shape of the mandibular cartilage altered in mutant animals. These Findings highlighted the role of low-frequency coding variants in normal-range facial variation.

## Introduction

Significant progress has been made in elucidating the genetic basis of human facial traits (Richmond, Howe, Lewis, Stergiakouli, & Zhurov, 2018; Weinberg, Cornell, & Leslie, 2018; Weinberg et al., 2019). Genome-wide association studies (GWASs) have identified and replicated numerous common genetic variants associated with normal-range facial morphology (Adhikari et al., 2016; Cha et al., 2018; Claes et al., 2018; Cole et al., 2016; Crouch et al., 2018; M. K. Lee et al., 2017; F. Liu et al., 2012; Paternoster et al., 2012; Shaffer et al., 2016); yet these variants cumulatively explain only a small fraction of the heritable phenotypic variation. Based on large-scale genomic studies of other complex morphological traits such as height (D. J. Liu et al., 2017; Lu et al., 2017; Marouli et al., 2017), we hypothesized that functional variants at hundreds or perhaps thousands of loci have yet to be discovered. While we expect that common variants, with a minor allele frequency (MAF) greater than 1%, account for most of the heritable variation in facial morphology, low frequency (MAF<1%) genetic variants may also play an important role. An exome-wide study of human height, for example, discovered 29 low-frequency coding variants with large effects of up to 2 centimeters per allele (Marouli et al., 2017).

Our previous GWAS in a modestly sized cohort of healthy individuals identified 1932 common genetic variants associated with facial variation at 38 loci, 15 of which were independently replicated (Claes et al., 2018). The success of this GWAS was attributed in part to an innovative data-driven phenotyping approach, in which the 3D facial surfaces were partitioned into hierarchically organized regions, each defined by multiple axes of shape variation. This approach allows for testing of genetic variants on facial morphology at multiple levels of scale – from the entire face (global) to highly localized facial regions (local). Extending this global-to-local analysis of facial traits to the analysis of low-frequency variants requires an appropriate and scalable statistical framework capable of accommodating the multivariate nature of the facial shape variables. A recently developed statistical approach, MultiSKAT (Dutta, Scott, Boehnke, & Lee, 2019), was designed for this purpose and showed desirable performance in its original development.

In this study, we evaluated the influence of low frequency coding variants, captured by the Illumina HumanExome BeadChip, on normal-range facial morphology in 2,329 individuals. We applied multivariate gene-based association testing methods to multi-dimensional facial shape phenotypes derived from 3D facial images. The results of our analyses pointed to novel genes, including at least one involved in orofacial clefts and several others with no previously described role in craniofacial development or disorders. We provided experimental evidence of our genetic association results through expression screening and knockout experiments in a zebrafish model. These results enhance our understanding of the genetic architecture of human facial variation.

## Materials and Methods

### Ethics statement

Institutional ethics (IRB) approval was obtained at each recruitment site and all subjects gave their written informed consent prior to participation (University of Pittsburgh Institutional Review Board #PRO09060553 and #RB0405013; UT Health Committee for the Protection of Human Subjects #HSC-DB-09-0508; Seattle Children’s Institutional Review Board #12107; University of Iowa Human Subjects Office/Institutional Review Board #200912764 and #200710721).

### Sample and Phenotyping

The study cohort comprised 2,329 unrelated, healthy individuals of European ancestry aged 3 to 40 years. Participants were eligible if they had not experienced facial trauma, major surgery, congenital facial anomalies that could potentially affect their natural facial structure. 3D images of each participant’s resting face were captured via digital stereophotogrammetry using the 3dMD face camera system. The data-driven phenotyping approached has been described in detail in a previous work (Claes et al., 2018). Briefly, approximately 10,000 points—“quasi-landmarks”—were automatically placed across the facial surface, by a non-rigid registration of a standard facial template onto each surface. The result is that each quasi-landmark represents the same facial position across all participants (White et al., 2019). The configurations were then co-aligned to their mean using generalized Procrustes analysis (GPA). The quasi-landmarks were then clustered into groups of co-varying component points in order to partition the full face into two segments. GPA was repeated within each of the two segments, and the process was continued for a total of four iterations to generate a hierarchy of 31 facial segments (which we call modules) comprising overlapping groups of quasi-landmarks. The hierarchical structure is illustrated in Fig 1, where modules formed successive levels representing the shift from more globally integrated to more locally focused morphology. Shape variation within each module was represented by the 3D coordinates of all quasi-landmarks contained therein. To reduce the dimensionality, principal components analysis and parallel analysis were performed on the quasi-landmarks. The result was a set of 31 multivariate phenotypes made up of 8 to 50 principal components (PCs) that jointly captured near complete shape variance. The effects of sex, age, height, weight, facial size and genetic ancestry were corrected for at the phenotyping stage. These facial module phenotypes were successfully used in our previous GWAS of common variants (Claes et al., 2018), which demonstrated a clear advantage of this data-driven multivariate modeling approach for gene-mapping studies over the traditional utilization of *a priori* (Shaffer et al., 2016) and univariate (M. K. Lee et al., 2017) facial traits.

**Fig 1.**
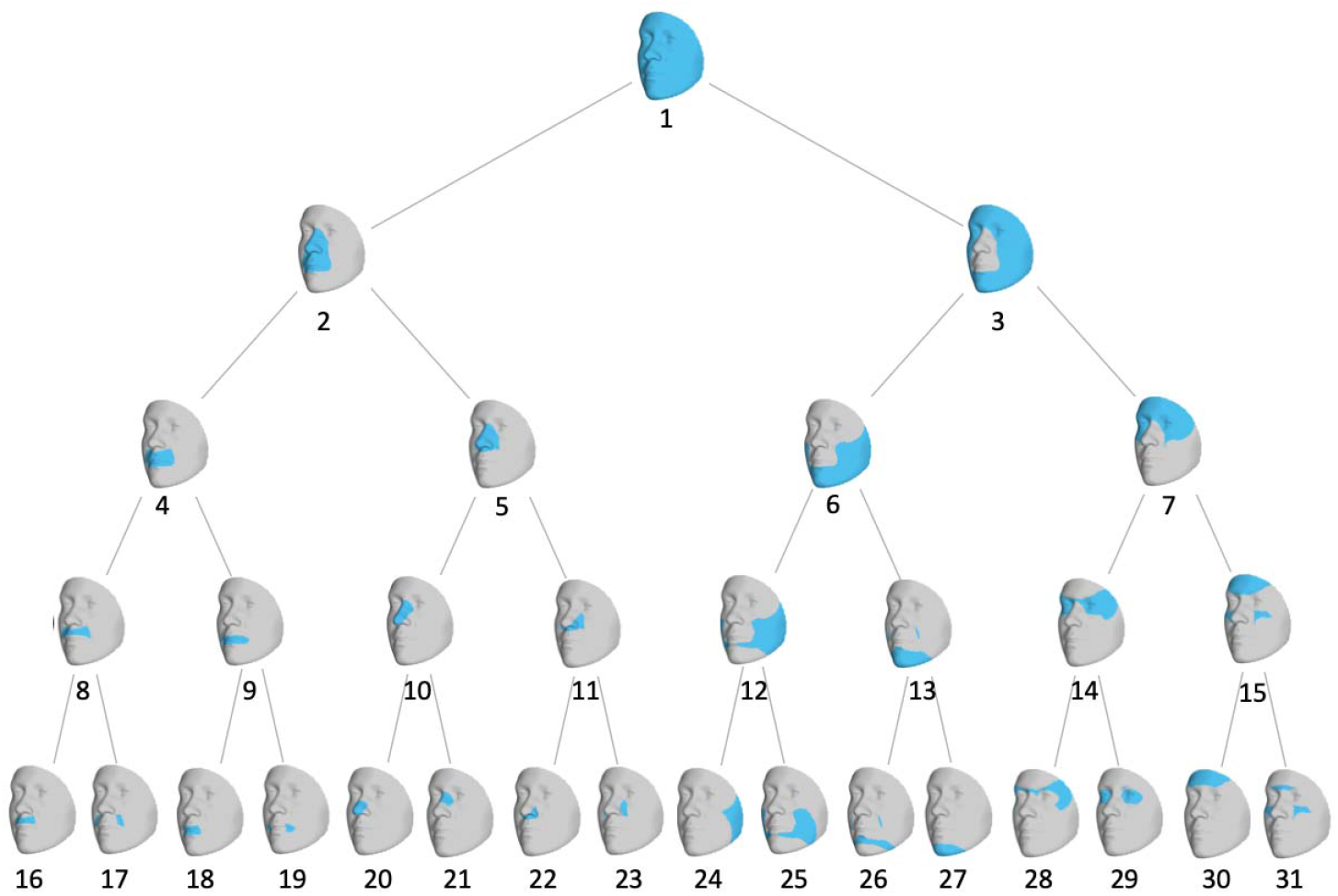
Hierarchical clustering of facial shape. Global-to-local facial segmentation obtained using hierarchical spectral clustering. Segments are colored in blue. The highest-level segment representing the full face was split into two sub-segments, and this bifurcation process was repeated until a five-level hierarchy comprising 31 segments was formed.

In addition to the phenotype quality control process described in (Claes et al., 2018), we further examined the phenotypic distribution of each module for extreme outlier faces, as phenotypic outliers may adversely impact low-frequency variant tests (Auer, Reiner, & Leal, 2016). To accomplish this, we looked at both the joint and the pairwise distribution of all PCs underlying each module. We visualized quantile-quantile (Q-Q) plots of chi-squared quantiles versus robust squared Mahalanobis distances to identify outliers that deviated from the rest of the sample. Mahalanobis distance is a metric measuring how far an observation is to the center of the joint distribution (centroid equivalent in a multivariate space). We identified one individual who was an outlier for several PCs in module 27 (chin), and revisited the associated facial images to confirm data validity and sample eligibility. This individual was excluded from any subsequent analysis involving module 27.

### Genotyping

Participants were genotyped by the Illumina OmniExpress + Exome v1.2 array, which included approximately 245,000 coding variants in the exome panel. Standard data cleaning and imputation procedures were implemented. Imputed genotypes with a certainty above 0.9 were used to fill in any sporadic missingness among genotype calls of the directly genotyped variants. We did not include any wholly unobserved, imputed SNPs in this analysis. Ancestry PCs based on common LD-pruned SNPs were constructed and regressed out from the multivariate traits to adjust for population structure.

### MultiSKAT

MultiSKAT (Dutta et al., 2019) was specifically developed for testing sets of variants, in this case coding variants within genes, for association with a multivariate trait. Testing low-frequency variants in aggregate can improve power compared to individual tests of each variant. The tool is flexible in relating multiple variants collectively to multiple phenotypes through the use of several choices of kernels, and includes an omnibus test to obtain optimal association p-values by integrating results across different kernels via Copula. This capability of accommodating multivariate phenotypes fits well with our analysis of facial modules, as each module was composed of several independent PCs. MultiSKAT can be applied to both common and rare variants, although our analysis considered low-frequency variants exclusively.

MultiSKAT uses a phenotype kernel to model how one variant affects multiple traits, and a separate genotype kernel to specify how multiple variants influence one trait. In reality, these effects are often not known *a priori*, and the true relationship can be a mixture of different effects. We used the heterogeneous and homogeneous phenotype kernels, which are appropriate when the set of traits analyzed are orthogonal PCs. We used the Sequence Kernel Association Test (SKAT) and burden test as the genotype kernel, and performed the omnibus test in MultiSKAT to aggregate results across the 2 × 2 kernel combinations.

### Gene-level analysis

Genome-wide coding variants with MAF<1% were aggregated into genes. Per the developer’s suggested practice for using the MultiSKAT method, we filtered out variants with three or fewer minor alleles to ensure that there is no inflation in MultiSKAT test statistics. We excluded genes with less than two qualified variants, leading to 31347 variants in 8091 genes being tested. When grouping multiple variants into a gene, MultiSKAT assigns larger weights to rarer variants. We applied a Bonferroni threshold to declare significance. To account for the correlation among partially overlapping facial modules, we used a procedure based on eigenvalues as proposed by Li and Ji (Li & Ji, 2005) and computed that the effective number of independent modules was 19. The threshold for significance was therefore set as p < 3.3 × 10^−7^ (i.e., 0.05/(8091 × 19)). The phenotypic effects of identified genes on face were visualized by creating and comparing the average facial morphs in individuals who had variants in a certain gene and those who do not carry any variants.

Gene-set enrichment analysis was carried out using GREAT (McLean et al., 2010), FUMA (Watanabe, Taskesen, van Bochoven, & Posthuma, 2017) and ToppFun (Chen, Bardes, Aronow, & Jegga, 2009). Expression of genes were looked up in the GTEx database (GTEx Consortium, 2013). Following our hypothesis that genes influencing typical facial presentation may also be involved in facial anomalies, we examined whether any genes identified by MultiSKAT were associated with non-syndromic cleft palate with or without cleft lip (NSCL/P) by retrieving association p-values from a past study of our group, where we performed a gene-based low-frequency variant association scan on NSCL/P (Leslie et al., 2017).

### Variant-level analysis

For genes highlighted by MultiSKAT, we scrutinized the quality of genotype calls by inspecting allele intensity cluster plots. We further performed association tests of individual SNPs using MultiPhen (O’Reilly et al., 2012). MultiPhen works by finding the linear combination of PCs that is mostly associated with the genotypes at each SNP, and is robust when variants with low frequencies are tested against non-normal phenotypes. Variant level functional prediction was performed using CADD (Rentzsch, Witten, Cooper, Shendure, & Kircher, 2019). CADD is a comprehensive metric that weights and integrates diverse sources of annotation, by contrasting variants that survived natural selection with simulated mutations. The scaled CADD score expresses the deleteriousness rank in terms of order of magnitude. A score of 10, for instance, is interpreted as ranking in the top 10% in terms of the damaging degree amongst reference genome SNPs, and a score of 20 refers to 1%, 30 to 0.1%, etc. Variant identifiers and chromosomal locations are indicated according to the hg19 genome build. Individual variants were searched in literature and PhenoScanner (Kamat et al., 2019) for existing human phenotype associations.

We quantified the magnitude of phenotypic effect of individual low-frequency variants by the difference between averaged faces of variant carriers (those who were heterozygotes; there was no homozygotes for the low-frequency variants tested) and non-carries, which was further compared with the effects of significant common variants identified in the prior GWAS of the same multidimensional traits (Claes et al., 2018). Specifically, the centroids of the multidimensional space defined by PCs in a certain module were computed separately for people carrying the variant and people who do not carry the variant. Then the Euclidean distance between the two centroids was calculated as a measure of variant effect size.

### Expression screen of candidate genes in zebrafish

The whole-mount RNA in situ hybridization (WISH) for *ar, cars2, ftsj1, hfe, ltb4r, telo2, nectin1a* and *nectin1b* was performed on wild type zebrafish embryos at 24 hpf and 48 hpf as described by Thisse et al. (C. Thisse & Thisse, 2008). All wild type embryos were collected synchronously at the corresponding stages and fixed in 4% paraformaldehyde (PFA) overnight. T7 RNA polymerase promoter was added to the reverse primers and was synthesized with antisense DIG-labeled probe in order to generate antisense RNA probe. The probe primers for *ar* are: forward 5’-GTCCTACAAGAACGCCAACG-3’ and reverse 5’-GGTCACAGACTTGGAAAGGG-3’ at 59°C. The *cars2* probe primers are: forward 5’-ATCTGGGTCATGCGTGTTCA-3’ and reverse 5’-GGATTCCTGTGGTGCTTGGT at 59°C. The *ftsj1* probe primers are: forward 5’-GGCGAGAAGTGCCTTCAAAC-3’ and reverse 5’-AGTCGTGCTTGTGTCTGGTT-3’ and *hfe* probe primers are: forward 5’-GGGGATGGATGCTTCTACGA-3’ and reverse 5’-CGCGCACACAAAATCATCAC-3’ at 59°C. The *ltb4r* probe primers are: forward 5’-GACGGTGCATTACCTGTGC-3’ and reverse 5’-AGTCTTGTCCGCCAAGGTC-3’ at 58°C. The primers for *telo2* are: forward 5’-GCTCCACTGGTGAGAGTGAG-3’ and reverse 5’-GTCAGCTGAGGAGAGTCTGCG-3’. The primers for *nectin1a* probe are: forward 5’-AACACCCAGGAGATCAGCAA-3’ and reverse 5’-CCTCCACCTCAGATCCGTAC-3’ at 57°C and the *nectin1b* probe primers are: forward 5’-TGCTAACCCAGCATTGGGAG-3’ and reverse 5’-GGTTCTTGGGCATTGGAGGA-3’ at 59°C. Embryos were mounted using glycerol and imaged using Nikon AZ100 multizoom microscope.

### Phenotype of mutant zebrafish

Zebrafish adults and embryos were obtained and maintained as described by Kimmel et al. (C. B. Kimmel, Ballard, Kimmel, Ullmann, & Schilling, 1995). Zebrafish nectin1a mutants were generated by transgene insertion Tg(Nlacz-GTvirus) in Chr 21: 21731876 - 21731886 (Zv9), and obtained from Zebrafish International Resource Center, allele Ia021885Tg (ZIRC catalog ID: ZL6899.07). The retroviral-mediated insertional mutagenesis inserts a molecular tag in the DNA and isolates the allele of interest. Therefore, this will induce a frameshift and probably causing either nonsense-mediated mRNA decay or a truncated protein (Amsterdam & Hopkins, 2003; Sivasubbu, Balciunas, Amsterdam, & Ekker, 2007). The PCR genotyping primers for *nectin1a* are: forward 5’-TTAGACCAGCCCACCTCA-3’ and reverse 5’-AATATGAAATAGCGCCGTTGTG-3’ at 62°C.

Alcian blue staining was performed as described by Walker et al. (Walker & Kimmel, 2007). The craniofacial cartilages were dissected and flat-mounted and then imaged using Nikon AZ100 multizoom microscope. After imaging, each embryo tail was placed in a PCR tube for genotyping. The protocol was used as described by (Westerfield, 1994) with modification of using fresh embryos without fixation.

## Results

In the gene-based test of exome-wide low-frequency variants, seven genes were significantly associated with one or more facial modules (*HFE, NECTIN1, CARS2, LTB4R, TELO2, AR*, and *FTSJ1;* Fig 2 and Table 1). Three of them showed associations with more than one module. Fig 3 and S1 Table show the results of these genes in multiple modules. Fig 3 shows the association signals propagating along the branching paths from the more global segments to the more local segments. Four genes (*HFE, CARS2, LTB4R*, and *TELO2*) were associated with nose-related modules, and the others were associated with the shape of chin, mouth, and cheek. *FTSJ1* had broad associations in the full face as well as in local regions, while the effects of other genes were more confined to only local modules. We observed well-calibrated test statistics and little evidence of inflation as shown in the Q-Q plots (Fig S1).

**Table 1.**
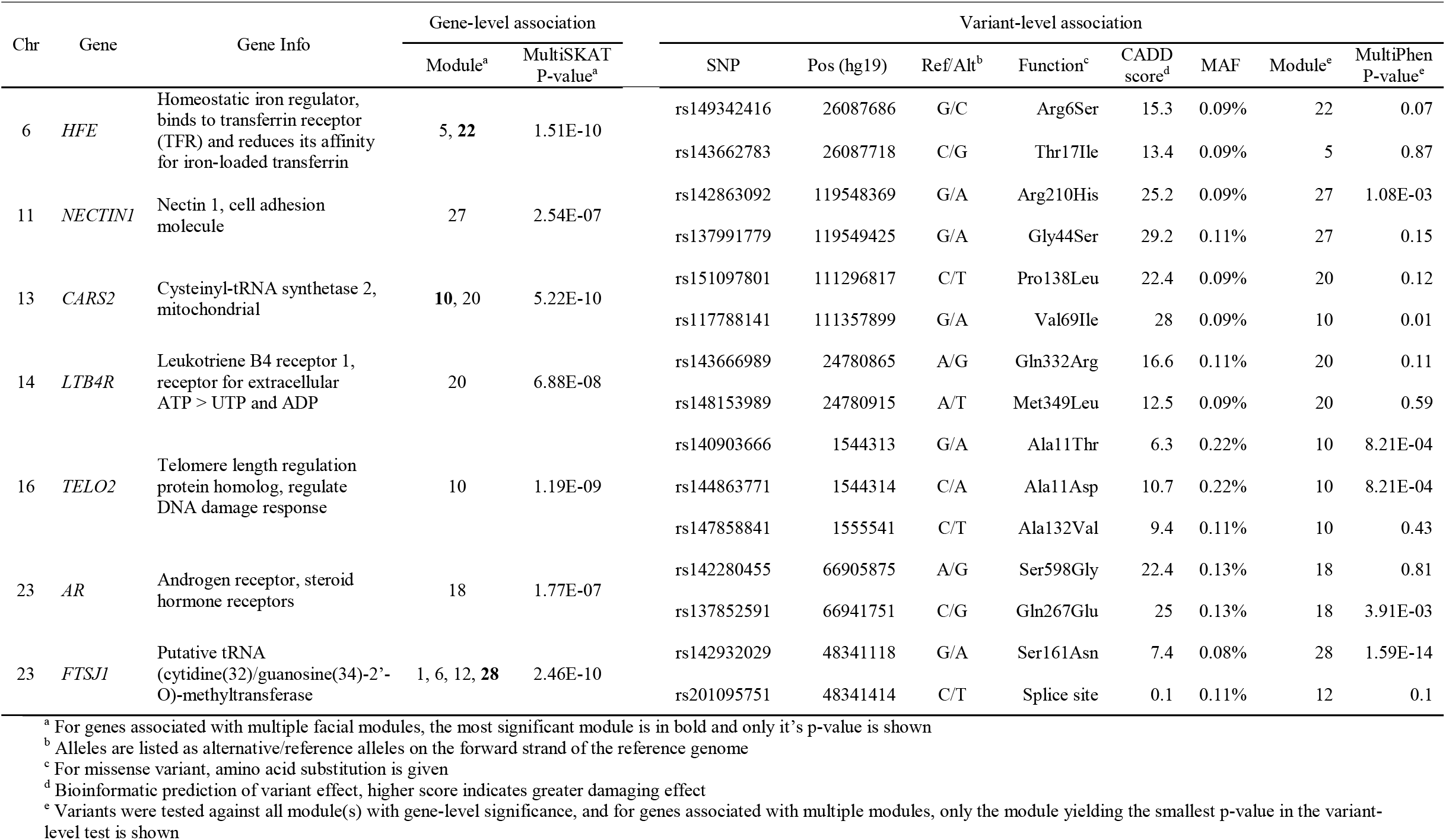
Single variant association and functional prediction for variants contributing to the gene-level significance

**Fig 2.**
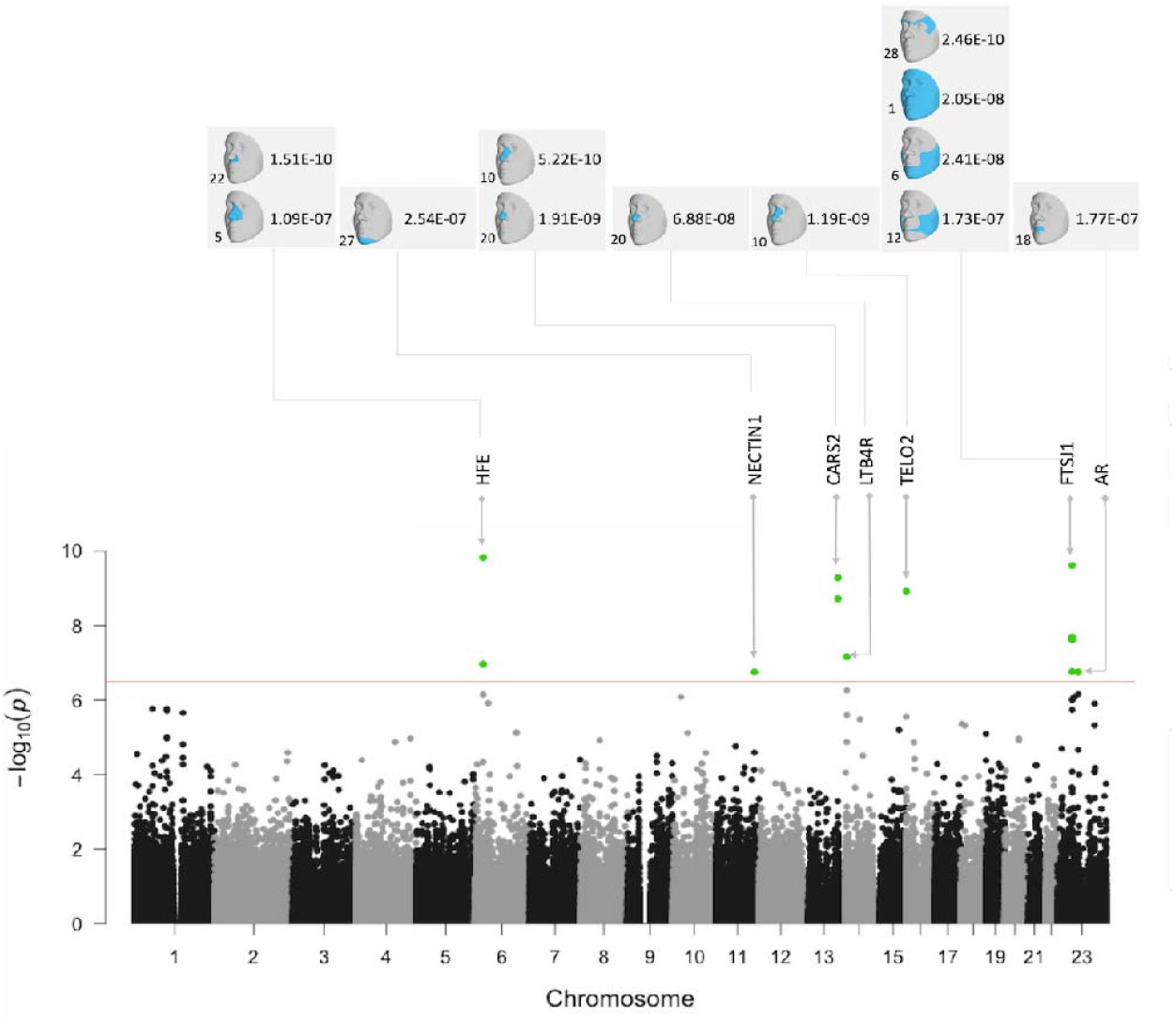
Composite Manhattan plot showing results across 31 facial modules. Manhattan plot showing the position of genes on the x axis and MultiSKAT p-values on the y axis. A total of 31 points are plotted for each gene, representing their p-values in each of the 31 modules. The red horizontal line indicates the significance threshold (3.3 × 10^−7^). The associated facial modules and the corresponding p-value for each gene that surpassed the threshold are shown above the Manhattan plot. The numbers to the bottom left of the facial images indicate the module identifiers in Fig 1.

**Fig 3.**
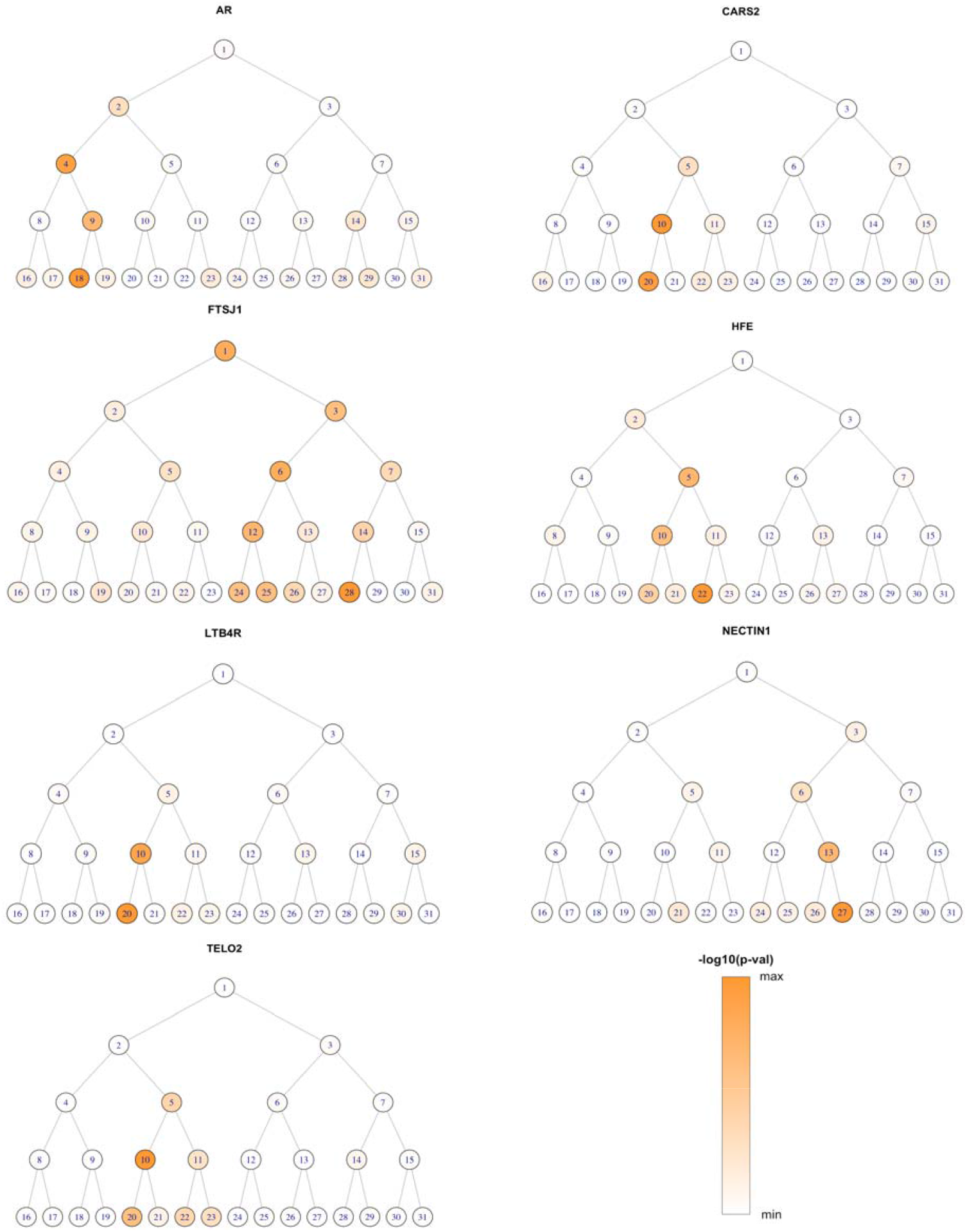
Module-wide association results for significant genes. For each gene, the –log10 p*-*value is shown as color shades ranging from min to max, for 31 facial segments arranged the same way as Fig 1. The global-to-local phenotyping enabled the discovery of genetic effects at different scales.

To visualize the effects of these genes on facial shape, we created the average module shape in non-carriers of the low-frequency variants for each gene, and a corresponding morph showing the change in shape from non-carriers to carriers (Fig 4). Blue and red indicate a local shape depression and protrusion, respectively, due to carrying any low-frequency variants. As an example, panel B in Fig 4 shows that *NECTIN1* variants shape the chin into a sharper and more protruding structure.

**Fig 4.**
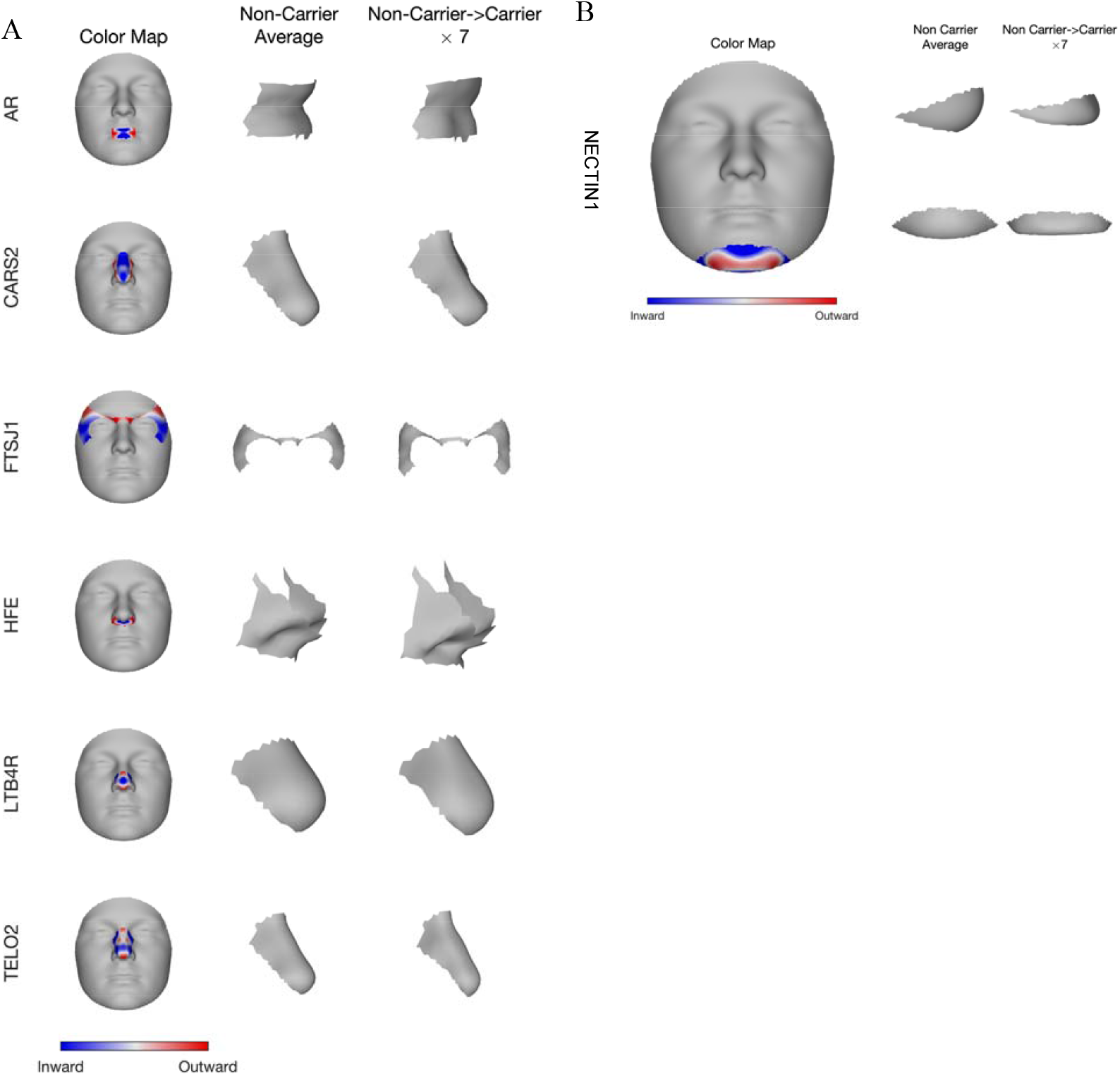
Phenotypic effect of the seven identified genes in their top associated module. Blue and red indicate a local shape depression and protrusion, respectively, due to carrying the low-frequency variants in the gene. A) First column shows gene effect on a representative module placing on the full face; middle column shows the lateral view of the average shape of the corresponding module among people who do not carry any variant in the gene; right column shows the change in the shape of the same module, from non-carrier to carrier, multiplied by a constant (7), to make the changes more visibly distinctive. B) For *NECTIN1* gene, we show both lateral (top) and frontal (bottom) view of its effect on chin shape. *NECTIN1* variant carriers on average displayed a sharper, more protruding chin.

We employed various bioinformatics tools to explore the functions associated with the set of identified genes. Enrichment was detected for a variety of biological processes (Fig S2), especially ion-, metabolism-, transport- and regulation-related processes. Enriched gene ontology (GO) molecular functions included signaling receptor and protein binding activity. Two genes with relatively well characterized functions (*HFE* and *AR)* contributed a lot to these enrichment results. In the GTEx database, these seven genes showed measurable expression level in adipose, skin and muscle-skeletal tissue (Fig S3), among which the strongest expression was seen for *NECTIN1* in skin.

To explore whether facial genes also affect the risk of orofacial clefts, results of gene-based associations of low-frequency (MAF<1%) variants with NSCL/P were retrieved from Leslie et al. 2017. Two out of the seven were not available from that study. S2 Table shows the SKAT and CMC test results for the other five genes in the European, Asian, South American and the combined samples. Two associations passed a Bonferroni corrected threshold for 40 tests (5 genes × 4 populations × 2 type of tests)— *TELO2* with a CMC p-value = 6.5 × 10^−4^, and HFE with a CMC p-value = 1.1 × 10^−3^, both in the combined population of all ancestry groups.

Single variants in the genes showing significant associations in the gene-based tests were further tested individually with the corresponding facial modules (Table 1). Six SNPs showed nominal associations (p-value < 0.05) and the top association involved SNP rs142932029 in *FTSJ1* with module 28 (p-value = 1.59 × 10^−14^). As shown in S4 Fig, these low-frequency variants had larger effects compared to previously reported common variants (Claes et al., 2018).

Most of the individual variants appeared at frequencies much lower than 1%, and all encode nonsynonymous substitutions except one splice site SNP in *FTSJ1*. Variants in *NECTIN1, CARS2* and *AR* are predicted to be deleterious according to their CADD score (details in S3 Table). SNP rs137991779 in *NECTIN1* has a CADD score of 29.2, interpreted as ranking in the top 0.12% in terms of deleteriousness among variants across the whole genome. PhenoScanner linked those variants with a variety of human traits/disorders in previous studies (S4 Table, mostly from UK Biobank), including height, vascular diseases, osteoporosis, neoplasms etc., suggesting that coding variants influencing facial shape may be pleiotropic and play roles in other biological processes.

Zebrafish WISH was used to examine *ar, cars2, ftsj1, hfe, ltb4r, telo2, nectin1a* and *nectin1b* expression pattern in the craniofacial region across key developmental stages (Fig 5). At 24 hours post fertilization (hpf), *ftsj1* was expressed in the hindbrain, and *hfe* and *ltb4r* were expressed in the forebrain. We detected nectin1a and nectin1b transcripts in the eyes, diencephalon, midbrain and hindbrain at 24 hpf. At 48 hpf, *ar* expression was detected in the epiphysis, *cars2, nectin1a* and *nectin1b* were expressed in the palate (Fig 5 solid arrow), and *nectin1a* was detected in the lower jaw (Fig 5 hollow arrow).

**Fig 5.**
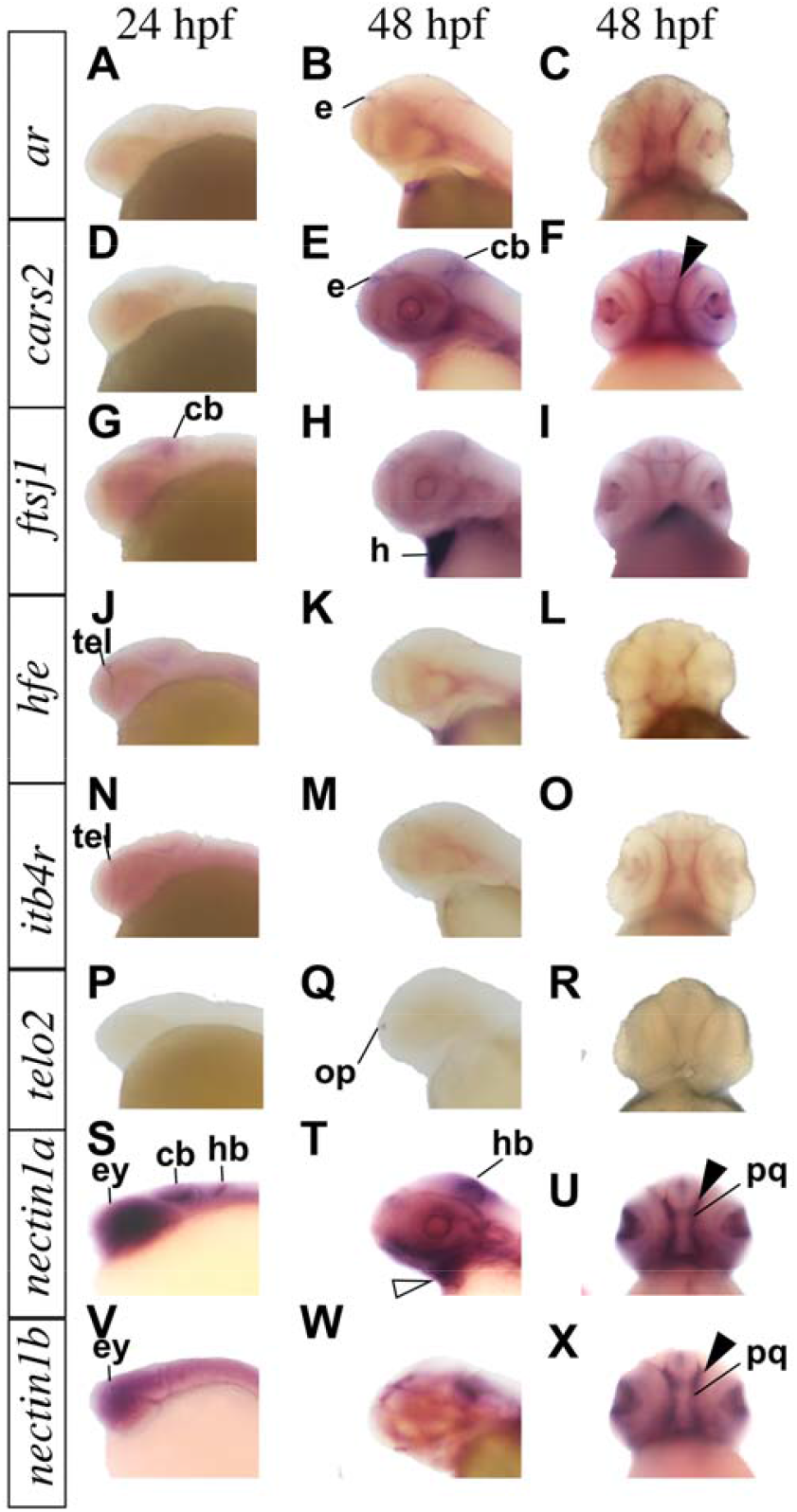
Whole-mount RNA in situ hybridization demonstrating genes expression in zebrafish. Genes expression pattern in lateral and ventral views at the indicated embryonic stages as hours per fertilization (hpf). *cars2, nectin1a* and *nectin1b* are expressed in zebrafish palate (solid arrow). *nectin1a* is expressed in the lower jaw at 48 hpf (hollow arrow). cb: cerebellum, e: epiphysis, ey: eye, h: heart, hb: hindbrain, op: olfactory placode, pq: palate quadrate, tel: telencephalon.

To investigate if *nectin1a* is required for during normal craniofacial development, we analyzed the *nectin1a* mutant allele Ia021885Tg. Breeding of *nectin1a*+/-intercross generated embryos with Mendelian ratio (1 individual homozygous for the wild type allele: 2 heterozygous individuals with one wild type and one mutant allele: 1 individual homozygous for the mutant allele) demonstrating a mutant craniofacial phenotype, characterized by small head structures (Fig 6). Using Alcain blue staining at 120 hpf, *nectin1a* mutants displayed dysmorphic craniofacial development with smaller and distorted palate and abnormal Meckel’s cartilage compared to age-matched wild type zebrafish embryos from the same intercross. These results show that *nectin1a* is genetically required for palate and mandible morphogenesis.

**Fig 6.**
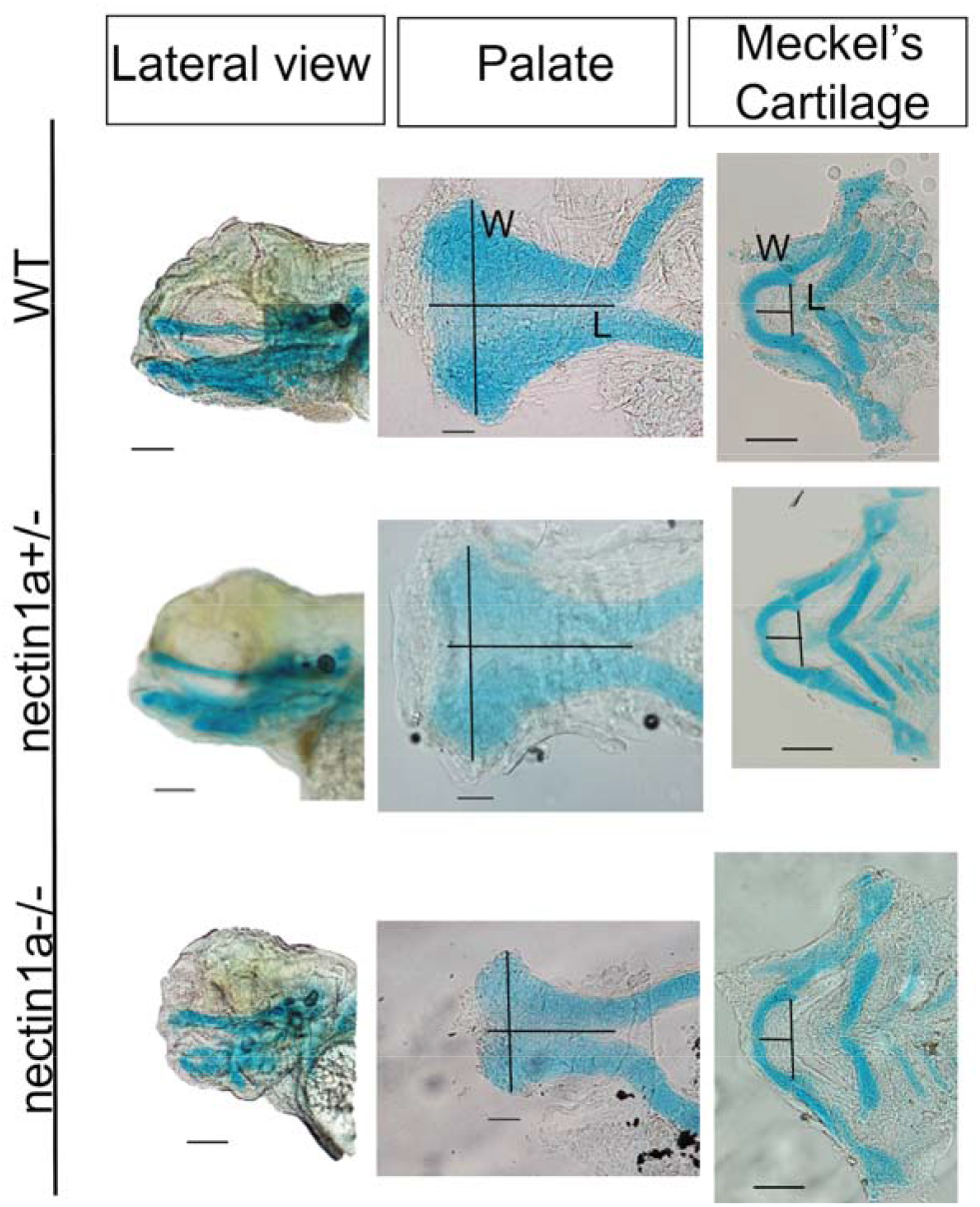
Alcian blue images for *nectin1a* zebrafish mutant comparted to wild type at day 5. Top images: wild type alcian blue lateral view, palate and Meckel’s cartilage. Middle images: heterozygous *nectin1a* embryo alcian blue. Bottom images: homozygous *nectin1a* mutant lateral view. The length of the palate was measured from the anterior midpoint to the posterior midpoint of the palate. The width was measured as the maximum distance between the 2 lateral borders at the anterior area. The length of the Meckel’s cartilage was measured from the midline of the Meckel’s cartilage to the midline of an imaginary line drawn joining the joints between the Meckel’s cartilage and the palatoquadrate. The width was measured from the junction of the Meckel’s cartilage and the palatoquadrate of one side to the other side. Compared to wild type animals. *nectin1a* mutants have smaller and shorter palate, and shorter and wider Meckel’s cartilage. L: length, W: width. Scale bar: 10 μm

## Discussion

This study presented a discovery effort to identify low-frequency coding variants associated with normal-range human facial shape, by undertaking gene-based association tests on a carefully phenotyped human cohort followed by functional experiments of the association results. Overall, we demonstrated that part of the morphological variation of facial shape is attributable to low-frequency coding variants, and pinpointed putative functional genes involved. Seven genes (*AR, CARS2, FTSJ1, HFE, LTB4R, TELO2* and *NECTIN1*) were identified, with phenotypic effects in the area of cheek, chin, nose and mouth. Notably, *NECTIN1* is known to cause a syndrome characterized by facial dysmorphology. Using a zebrafish model, we confirmed the expression of *nectin1a* and *nectin1b* in the developing head and the abnormal craniofacial phenotype in *nectin1a* mutants, with the affected structures being highly consistent with the associated facial region in the human data analysis. Taken together, these findings support the contribution of low-frequency coding variants to the genetic architecture of normal-range facial shape.

The seven genes identified by the multivariate approach are for the first time implicated in normal facial morphology. Six of the seven genes (all but *cars2)* were expressed in embryonic craniofacial tissues in zebrafish, demonstrating their potential involvement in craniofacial development. Cellular processes/functions of these genes include metal ion transport (*HFE*), signaling (*AR, LTB4R*), tRNA metabolism (*CARS2, FTSJ1*), DNA repair (*TELO2*) and cell adhesion (*NECTIN1*). This diversity in their biological function led to a variety of enriched functional pathways/categories in the gene-set enrichment analysis, yet without a strong signal in any particular one, probably due to the small number of genes and the polygenic nature of facial morphology. With the exception of *NECTIN1*, the role of these genes in patterning craniofacial structures is unknown, and further investigation is needed to gain better understanding of how these genes may affect the face.

Previous GWASs and studies of facial dysmorphology have demonstrated that there are common genetic factors underlying normal-range facial variation and orofacial clefting (Claes et al., 2018; F. Liu et al., 2012; Weinberg et al., 2009). Our findings suggest that low-frequency coding variants may also help explain this relationship. Although none of the other genes implicated here have been shown to be involved in craniofacial development, *NECTIN1* is an established player that has been linked to both syndromic and isolated forms of orofacial clefting (Avila et al., 2006; Sozen et al., 2001; Suzuki et al., 2000). Individuals with cleft lip/palate-ectodermal dysplasia syndrome (OMIM:225060) have distinctive facial features including an underdeveloped lower jaw (Zlotogora, 1994), which is consistent with the facial segment (chin) where the *NECTIN1* association was observed. Although not passing the genome-wide threshold, *NECTIN1* also yielded some signals in modules representing the nose and cheek (Fig 3), additional facial regions affected in this syndrome. Different variants in *NECTIN1* are likely involved in normal-range variation and in craniofacial disorders, which may help explain apparent differences in phenotypic severity. NECTIN1 protein belongs to the subfamily of immunoglobulin-like adhesion molecules which are key components of cell adhesion junctions and play critical roles in the development of many tissues, including in the fusion of palatal shelves during palatogenesis (Cobourne, 2004). A handful of *NECTIN1* mutations that can potentially disrupt gene function have been documented in non-syndromic cleft patients (Oner & Tastan, 2016; Scapoli et al., 2006; Tongkobpetch, Suphapeetiporn, Siriwan, & Shotelersuk, 2008). In the current study, two coding variants in *NECTIN1* contributed to the gene-level significance, both predicted to be deleterious. We performed lookups of the face-associated genes in a previous exome scan of a NSCL/P cohort (Leslie et al., 2017). *NECTIN1* yielded a small p-value of 0.004, although not passing the Bonferroni significance threshold. Two other genes, *TELO2* and *HFE*, did pass that threshold. These results are in line with previous evidence suggesting a role for same genes in normal and abnormal facial development.

Our zebrafish experiments provided a strong support for the relevance of *nectin1a* in palate and mandible development. The mutants displayed changes in the shape and size of both the palate and the Meckel’s cartilage, from which the mandibles evolved. This affected cartilage structure in zebrafish mutants aligns well with the associated human anatomical region (chin and mandible), where the effects of *NECTIN1* were observed in the MultiSKAT test. These findings for the first time demonstrate a role of *NECTIN1* in normal-range facial variation. We highlight the approach of interrogating human candidate genes in a biological context using the zebrafish model, where dynamic gene expression can be assayed in a high throughput fashion. Those candidate genes with spatiotemporal gene expression in the craniofacial domains then can be evaluated in functional studies, were mutants may already be available from large scale mutagenesis projects, or can be generated by CRISPR mediated gene editing.

With the hierarchical facial segmentation, we were able to identify genetic effects at different scales. For example, the effects of *FTSJ1* were observed globally in the full face, and also locally in specific modules on the side of the face. By contrast, the effect of *NECTIN1* was confined to localized facial parts only. These patterns may help with understanding the mechanisms by which genes act along the growth of facial structure. Our multivariate data-driven phenotyping approach eliminates the need of preselecting traits, captures more variation in the facial shape, and is more effective for gene mapping.

The current study is an important extension and complement of our prior work on common SNPs (Claes et al., 2018). Here we exclusively focused on coding variants with MAF below 1%, which have been omitted based no standard QC procedure from previous facial GWAS attempts. We compared results from this study to those from our prior GWAS (Claes et al., 2018), and noted that common variants in or near (within 500kb) the seven associated genes showed no evidence of association (p > 0.001 for all) with the same facial modules. This indicates that the current study generated distinct, non-overlapping knowledge on facial genetics, although it is possible that there are trans-acting common GWAS SNPs that regulate the expression of the seven identified genes during facial morphogenesis. Low-frequency variants showed larger magnitude of effects compared to common variants in (Claes et al., 2018). It is necessary, however, to point out that this difference could partially or completely be a result of the drastically smaller groups of variant carriers, and we therefore refrain from overinterpreting the comparison.

Our study demonstrated the power of applying gene-based tests of low-frequency variants that are usually untestable individually. While some significant genes harbor variants with a small p-value in our single-variant association test, others would have been missed if not tested in aggregate. With a moderate sample size of 2329, it is highly desirable to collapse low-frequency variants into putative functional units and perform burden-style tests. In addition to an increase in power, another key benefit with analyzing low-frequency coding variants collectively is the improved biological interpretability compared to GWASs. The gene-centered design of coding variant tests facilities much clearer biological implications and options for experimental follow-up. Our success with the functional validation of *NECTIN1* provides a practical example. We expect future better-powered studies to discover more biological pathways emerging from analyses of low-frequency coding variants.

Replication of rare and low-frequency variant association signals presents unique challenges. The prominent barrier is the limited sample size. The low numbers or even absence of the carriers in independent populations hindered the replication efforts of our findings. Six out of the seven genes identified were not testable in a separate cohort of 664 participants due to a lack of variant carriers. Given our sample size and the ExomeChip design, this study was not adequately powered to identify genes harboring extra rare variants that may also contribute to facial traits. Although complex traits are not expected to have a large fraction of the heritability explained by rare and private variants, such variants may be influential, predictive, and actionable at the individual level. In this regard, whole exome or whole genome sequencing of large samples holds promise to give deeper insights into the role rare variants in facial morphology.

Like many other complex traits, research with a focus on uncovering the genetic architecture of facial morphology is confronted with the challenge of missing heritability (Cole et al., 2017; Tsagkrasoulis, Hysi, Spector, & Montana, 2017). Our study has extended the paradigm of genetic factors involved in facial morphology from common to low frequency variants and highlighted novel candidate genes that may lead to encouraging follow-ups. Given that rare and low-frequency genetic variation might be highly specific to certain populations, and facial shapes have distinctive ancestry features, future studies may benefit from extending the discovery of influential low-frequency variants to other ethnic groups.

## Acknowledgements

The authors thank all the dedicated staff, collaborators, and participants for their contribution to the study.

## Data availability

All of the genotypic markers are available to the research community through the dbGaP controlled-access repository (https://dbgap.ncbi.nlm.nih.gov/) at accession phs000949.v1.p1. The raw source data for the phenotypes – the 3D facial surface models – are available for the 3D Facial Norms dataset through the FaceBase Consortium (www.facebase.org). Access to these 3D facial surface models requires proper institutional ethics approval and approval from the FaceBase data access committee. KU Leuven provides the spatially dense facial mapping software, free to use for academic purposes: MeshMonk (https://github.com/TheWebMonks/meshmonk).

## Supporting Information

**S1 Fig. Q-Q plot of gene-based MultiSKAT tests by facial module**

**S2 Fig. FUMA enrichment results**

**S3 Fig. GTEx expression of MultiSKAT significant genes in tissues relevant to facial morphology**. Dendrogram denotes similarity in expression level. TPM, transcripts per million

**S4 Fig. Magnitude of variant effect on facial modules, quantified by the Euclidean distance between averaged faces of different genotype groups**. The 95% confidence interval was obtained by 5000 bootstraps. The farther away the blue (common) or red (low-freq) rectangular boxes fall from line x=0, the larger the group distances and the greater the magnitude of effects. Common variants that yielded significant GWAS association in the same cohort with the same modules are used as a comparison to low-frequency variants. Genotype groups column indicates the two groups of people of whom the faces were averaged and distance was computed. For example, 0 vs 1/2 means minor allele homozygotes vs the remaining. The following two columns indicate sizes of the two groups in comparison. Low-frequency variants had large effects compared to previously reported common variants, although this could be a result from the much smaller size of carrier group and may not reflect genuine greater effects of low-frequency variants.

**S1 Table. Module-wide association results of genes identified by MultiSKAT**. Show modules with a p-value < 10E-4.

**S2 Table. SKAT and CMC test results of the association between the seven facial genes and NSCL/P, retrieved from a previous exome-wide gene-based association study of NSCL/P**

**S3 Table. Functional prediction of individual variants in significant genes by CADD GRCh37-v1**.**4**

**S4 Table. PhenoScanner lookups for variants in seven significant genes**. Show existing associations involving these variants with a p value < 10e-4.

## Reference

Adhikari, K., Fuentes-Guajardo, M., Quinto-Sánchez, M., Mendoza-Revilla, J., Camilo Chacón-Duque, J., Acuña-Alonzo, V., et al. (2016). A genome-wide association scan implicates DCHS2, RUNX2, GLI3, PAX1 and EDAR in human facial variation. Nature Communications, 7(1), 11616. http://doi.org/10.1038/ncomms11616

Amsterdam, A., & Hopkins, N. (2003). Retroviral-Mediated Insertional Mutagenesis in Zebrafish. Methods in Cell Biology, 77, 3–20. http://doi.org/10.1016/S0091-679X(04)77001-6

Auer, P. L., Reiner, A. P., & Leal, S. M. (2016). The effect of phenotypic outliers and non-normality on rare-variant association testing. European Journal of Human Genetics : EJHG, 24(8), 1188–1194. http://doi.org/10.1038/ejhg.2015.270

Avila, J. R., Jezewski, P. A., Vieira, A. R., Orioli, I. M., Castilla, E. E., Christensen, K., et al. (2006). PVRL1 variants contribute to non-syndromic cleft lip and palate in multiple populations. American Journal of Medical Genetics Part A, 140(23), 2562–2570. http://doi.org/10.1002/ajmg.a.31367

Cha, S., Lim, J. E., Park, A. Y., Do, J.-H., Lee, S. W., Shin, C., et al. (2018). Identification of five novel genetic loci related to facial morphology by genome-wide association studies. Bmc Genomics, 19(1), 481–17. http://doi.org/10.1186/s12864-018-4865-9

Chen, J., Bardes, E. E., Aronow, B. J., & Jegga, A. G. (2009). ToppGene Suite for gene list enrichment analysis and candidate gene prioritization. Nucleic Acids Research, 37(Web Server), W305–W311. http://doi.org/10.1093/nar/gkp427

Claes, P., Roosenboom, J., White, J. D., Swigut, T., Sero, D., Li, J., et al. (2018). Genome-wide mapping of global-to-local genetic effects on human facial shape. Nature Genetics, 50(3), 1–16. http://doi.org/10.1038/s41588-018-0057-4

Cobourne, M. T. (2004). The complex genetics of cleft lip and palate. European Journal of Orthodontics, 26(1), 7–16.

Cole, J. B., Manyama, M., Kimwaga, E., Mathayo, J., Larson, J. R., Liberton, D. K., et al. (2016). Genomewide Association Study of African Children Identifies Association of SCHIP1 and PDE8A with Facial Size and Shape. PLoS Genetics, 12(8), e1006174. http://doi.org/10.1371/journal.pgen.1006174

Cole, J. B., Manyama, M., Larson, J. R., Liberton, D. K., Ferrara, T. M., Riccardi, S. L., et al. (2017). Human Facial Shape and Size Heritability and Genetic Correlations. Genetics, 205(2), 967–978. http://doi.org/10.1534/genetics.116.193185

Crouch, D. J. M., Winney, B., Koppen, W. P., Christmas, W. J., Hutnik, K., Day, T., et al. (2018). Genetics of the human face: Identification of large-effect single gene variants. Proceedings of the National Academy of Sciences of the United States of America, 115(4), E676–E685. http://doi.org/10.1073/pnas.1708207114

Dutta, D., Scott, L., Boehnke, M., & Lee, S. (2019). Multi-SKAT: General framework to test for rare-variant association with multiple phenotypes. Genetic Epidemiology, 43(1), 4–23. http://doi.org/10.1002/gepi.22156

GTEx Consortium. (2013). The Genotype-Tissue Expression (GTEx) project. Nature Genetics, 45(6), 580–585. http://doi.org/10.1038/ng.2653

Kamat, M. A., Blackshaw, J. A., Young, R., Surendran, P., Burgess, S., Danesh, J., et al. (2019). PhenoScanner V2: an expanded tool for searching human genotype-phenotype associations. Bioinformatics, 35(22), 4851–4853. http://doi.org/10.1093/bioinformatics/btz469

Kimmel, C. B., Ballard, W. W., Kimmel, S. R., Ullmann, B., & Schilling, T. F. (1995). Stages of Embryonic-Development of the Zebrafish. Developmental Dynamics, 203(3), 253–310. http://doi.org/10.1002/aja.1002030302

Lee, M. K., Shaffer, J. R., Leslie, E. J., Orlova, E., Carlson, J. C., Feingold, E., et al. (2017). Genome-wide association study of facial morphology reveals novel associations with FREM1 and PARK2. PLoS ONE, 12(4), e0176566–13. http://doi.org/10.1371/journal.pone.0176566

Leslie, E. J., Carlson, J. C., Shaffer, J. R., Buxó, C. J., Castilla, E. E., Christensen, K., et al. (2017). Association studies of low-frequency coding variants in nonsyndromic cleft lip with or without cleft palate. American Journal of Medical Genetics Part B: Neuropsychiatric Genetics, 173(6), 1531–1538. http://doi.org/10.1002/ajmg.a.38210

Li, J., & Ji, L. (2005). Adjusting multiple testing in multilocus analyses using the eigenvalues of a correlation matrix. Heredity, 95(3), 221–227. http://doi.org/10.1038/sj.hdy.6800717

Liu, D. J., Peloso, G. M., Yu, H., Butterworth, A. S., Wang, X., Mahajan, A., et al. (2017). Exome-wide association study of plasma lipids in >300,000 individuals. Nature Genetics, 49(12), 1758–1766. http://doi.org/10.1038/ng.3977

Liu, F., van der Lijn, F., Schurmann, C., Zhu, G., Chakravarty, M. M., Hysi, P. G., et al. (2012). A Genome-Wide Association Study Identifies Five Loci Influencing Facial Morphology in Europeans. PLoS Genetics, 8(9), e1002932–13. http://doi.org/10.1371/journal.pgen.1002932

Lu, X., Peloso, G. M., Liu, D. J., Wu, Y., Zhang, H., Zhou, W., et al. (2017). Exome chip meta-analysis identifies novel loci and East Asian–specific coding variants that contribute to lipid levels and coronary artery disease. Nature Genetics, 49(12), 1722–1730. http://doi.org/10.1038/ng.3978

Marouli, E., Graff, M., Medina-Gomez, C., Lo, K. S., Wood, A. R., Kjaer, T. R., et al. (2017). Rare and low-frequency coding variants alter human adult height. Nature, 542(7640), 186–190. http://doi.org/10.1038/nature21039

McLean, C. Y., Bristor, D., Hiller, M., Clarke, S. L., Schaar, B. T., Lowe, C. B., et al. (2010). GREAT improves functional interpretation of cis-regulatory regions. Nature Biotechnology, 28(5), 495–501. http://doi.org/10.1038/nbt.1630

Oner, D. A., & Tastan, H. (2016). Identification of Novel Variants in the PVRL1 Gene in Patients With Nonsyndromic Cleft Lip With or Without Cleft Palate. Genetic Testing and Molecular Biomarkers, 20(5), 269–272. http://doi.org/10.1089/gtmb.2015.0276

O’Reilly, P. F., Hoggart, C. J., Pomyen, Y., Calboli, F. C. F., Elliott, P., Jarvelin, M.-R., & Coin, L. J. M. (2012). MultiPhen: Joint Model of Multiple Phenotypes Can Increase Discovery in GWAS. PLoS ONE, 7(5), e34861–12. http://doi.org/10.1371/journal.pone.0034861

Paternoster, L., Zhurov, A. I., Toma, A. M., Kemp, J. P., Pourcain, B. S., Timpson, N. J., et al. (2012). Genome-wide Association Study of Three-Dimensional Facial Morphology Identifies a Variant in PAX3 Associated with Nasion Position. The American Journal of Human Genetics, 90(3), 478–485. http://doi.org/10.1016/j.ajhg.2011.12.021

Rentzsch, P., Witten, D., Cooper, G. M., Shendure, J., & Kircher, M. (2019). CADD: predicting the deleteriousness of variants throughout the human genome. Nucleic Acids Research, 47(D1), D886–D894. http://doi.org/10.1093/nar/gky1016

Richmond, S., Howe, L. J., Lewis, S., Stergiakouli, E., & Zhurov, A. (2018). Facial Genetics: A Brief Overview. Frontiers in Genetics, 9, 462. http://doi.org/10.3389/fgene.2018.00462

Scapoli, L., Palmieri, A., Martinelli, M., Vaccari, C., Marchesini, J., Pezzetti, F., et al. (2006). Study of the PVRL1 gene in Italian nonsyndromic cleft lip patients with or without cleft palate. Annals of Human Genetics, 70(Pt 3), 410–413. http://doi.org/10.1111/j.1529-8817.2005.00237.x

Shaffer, J. R., Orlova, E., Lee, M. K., Leslie, E. J., Raffensperger, Z. D., Heike, C. L., et al. (2016). Genome-Wide Association Study Reveals Multiple Loci Influencing Normal Human Facial Morphology. PLoS Genetics, 12(8), e1006149–21. http://doi.org/10.1371/journal.pgen.1006149

Sivasubbu, S., Balciunas, D., Amsterdam, A., & Ekker, S. C. (2007). Insertional mutagenesis strategies in zebrafish. Genome Biology, 8(S1), S9. http://doi.org/10.1186/gb-2007-8-s1-s9

Sozen, M. A., Suzuki, K., Tolarova, M. M., Bustos, T., Fernández Iglesias, J. E., & Spritz, R. A. (2001). Mutation of PVRL1 is associated with sporadic, non-syndromic cleft lip/palate in northern Venezuela. Nature Genetics, 29(2), 141–142. http://doi.org/10.1038/ng740

Suzuki, K., Hu, D., Bustos, T., Zlotogora, J., Richieri-Costa, A., Helms, J. A., & Spritz, R. A. (2000). Mutations of PVRL1, encoding a cell-cell adhesion molecule/herpesvirus receptor, in cleft lip/palate-ectodermal dysplasia. Nature Genetics, 25(4), 427–430. http://doi.org/10.1038/78119

Thisse, C., & Thisse, B. (2008). High-resolution in situ hybridization to whole-mount zebrafish embryos. Nature Protocols, 3(1), 59–69. http://doi.org/10.1038/nprot.2007.514

Tongkobpetch, S., Suphapeetiporn, K., Siriwan, P., & Shotelersuk, V. (2008). Study of the poliovirus receptor related-1 gene in Thai patients with non-syndromic cleft lip with or without cleft palate. International Journal of Oral & Maxillofacial Surgery, 37(6), 550–553. http://doi.org/10.1016/j.ijom.2008.01.024

Tsagkrasoulis, D., Hysi, P., Spector, T., & Montana, G. (2017). Heritability maps of human face morphology through large-scale automated three-dimensional phenotyping. Scientific Reports, 7(1), 1–18. http://doi.org/10.1038/srep45885

Walker, M. B., & Kimmel, C. B. (2007). A two-color acid-free cartilage and bone stain for zebrafish larvae. Biotechnic & Histochemistry, 82(1), 23–28. http://doi.org/10.1080/10520290701333558

Watanabe, K., Taskesen, E., van Bochoven, A., & Posthuma, D. (2017). Functional mapping and annotation of genetic associations with FUMA. Nature Communications, 8(1), 1826. http://doi.org/10.1038/s41467-017-01261-5

Weinberg, S. M., Cornell, R., & Leslie, E. J. (2018). Craniofacial genetics: Where have we been and where are we going? PLoS Genetics, 14(6), e1007438. http://doi.org/10.1371/journal.pgen.1007438

Weinberg, S. M., Naidoo, S. D., Bardi, K. M., Brandon, C. A., Neiswanger, K., Resick, J. M., et al. (2009). Face shape of unaffected parents with cleft affected offspring: combining three-dimensional surface imaging and geometric morphometrics. Orthodontics & Craniofacial Research, 12(4), 271–281. http://doi.org/10.1111/j.1601-6343.2009.01462.x

Weinberg, S. M., Roosenboom, J., Shaffer, J. R., Shriver, M. D., Wysocka, J., & Claes, P. (2019). Hunting for genes that shape human faces: Initial successes and challenges for the future. Orthodontics & Craniofacial Research, 22 Suppl 1(S1), 207–212. http://doi.org/10.1111/ocr.12268

Westerfield, M. (1994). A guide for the laboratory use of zebrafish Danio (Brachydanio) rerio. In: University of Oregon Press, Eugene.

White, J. D., Ortega-Castrillón, A., Matthews, H., Zaidi, A. A., Ekrami, O., Snyders, J., et al. (2019). MeshMonk: Open-source large-scale intensive 3D phenotyping. Scientific Reports, 9(1), 6085. http://doi.org/10.1038/s41598-019-42533-y

Zlotogora, J. (1994). Syndactyly, Ectodermal Dysplasia, and Cleft Lip/Palate. Journal of Medical Genetics, 31(12), 957–959. http://doi.org/10.1136/jmg.31.12.957

