## Supplementary figures and images for "Impact of low-frequency coding variants on human facial shape"

### Fig S1

Fig S1. Q-Q plot of gene-based MultiSKAT tests by facial module

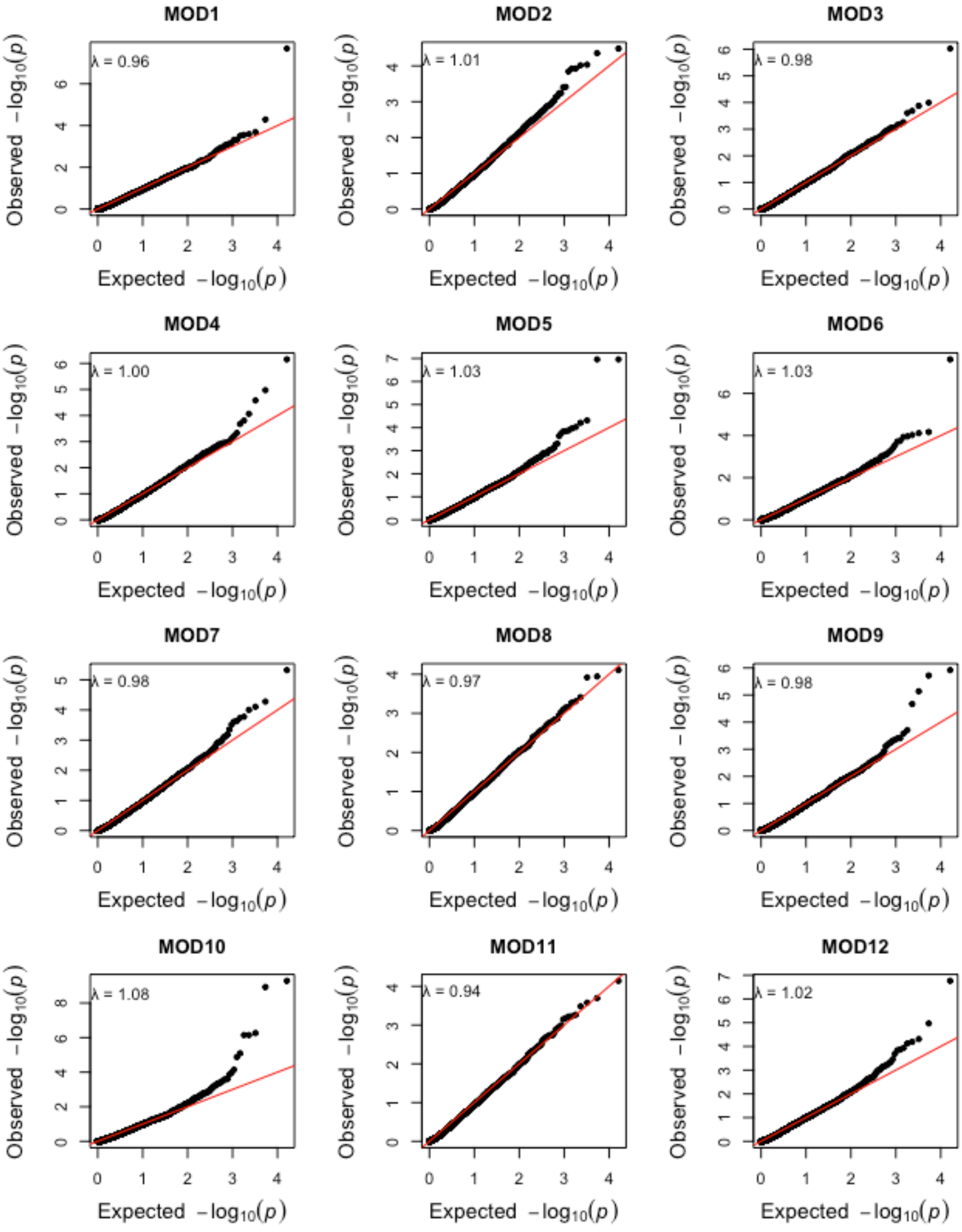

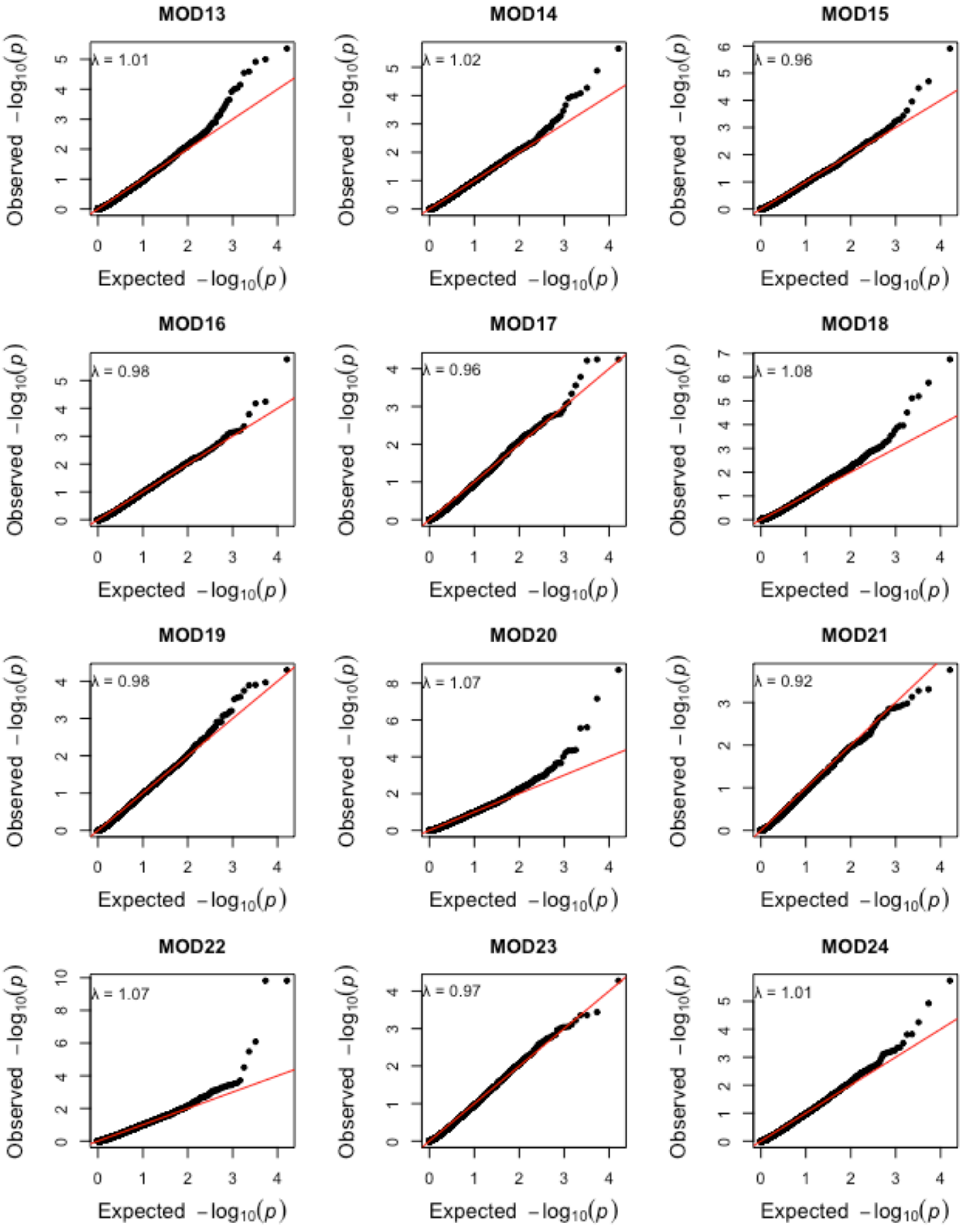

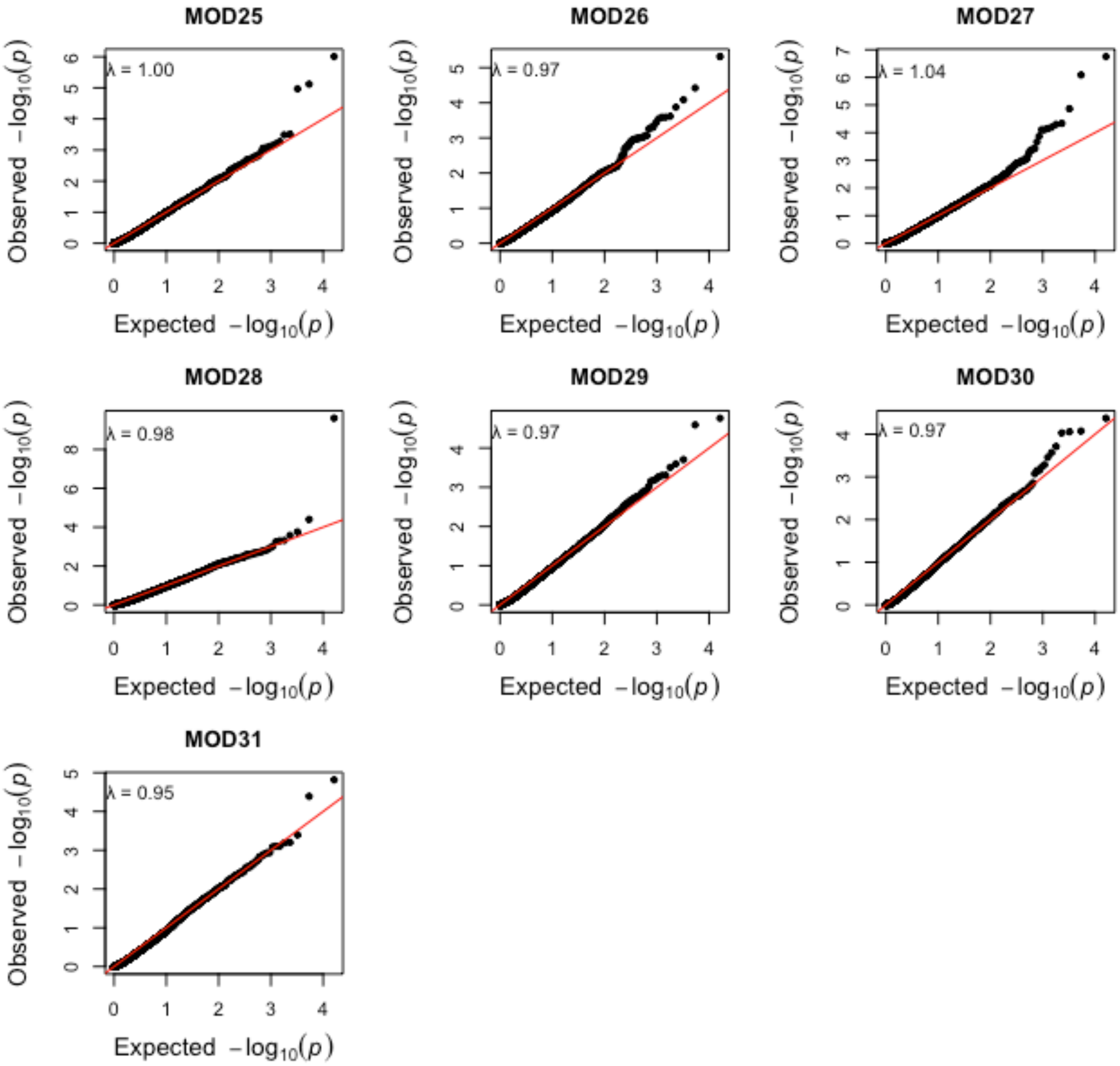

### Fig S2

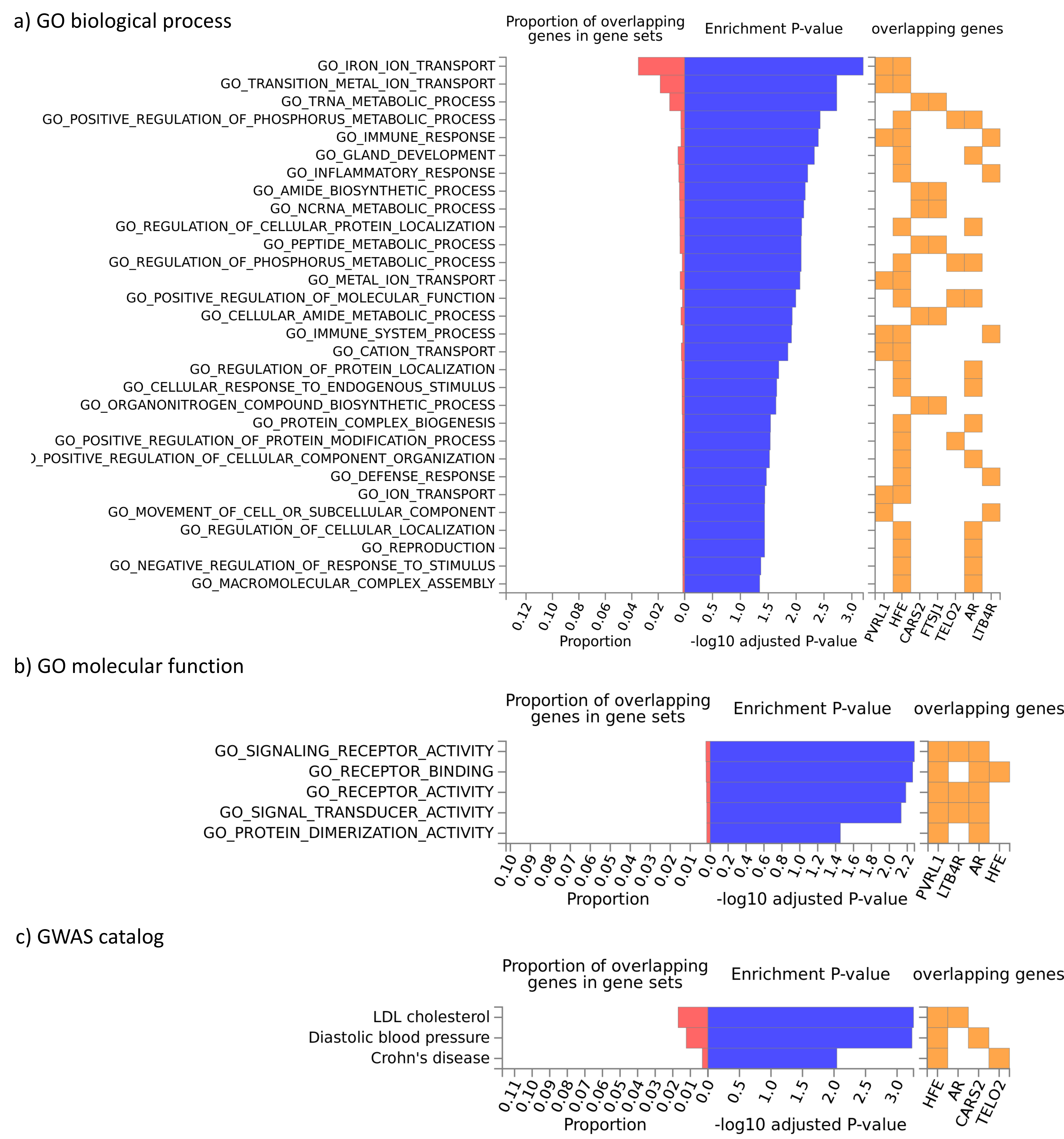
