## Supplementary material for "Impact of low-frequency coding variants on human facial shape": Fig S3

Fig S3. GTEx expression of MultiSKAT significant genes in tissues relevant to facial morphology. Dendrogram denotes similarity in expression level. TPM, transcripts per million.

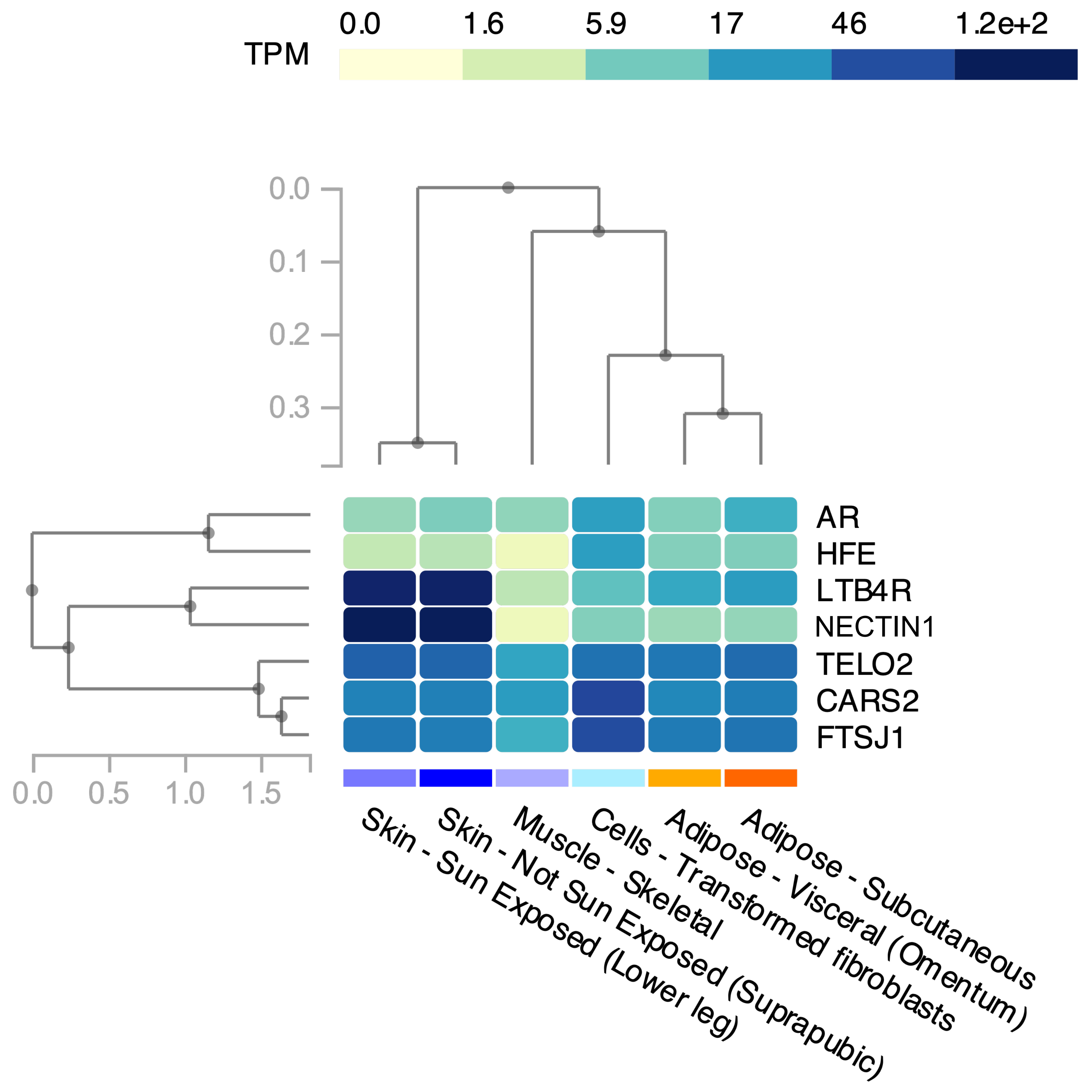
