## Supplementary material for "Impact of low-frequency coding variants on human facial shape": Fig S4

| Module | Variant | Genotype groups | N_noncarriers | N_carriers |
|---|---|---|---|---|
| **Common variants** | | | | |
| 10 | rs227833 | 0 vs 1 | 1266 | 909 |
| | | 0 vs 2 | 1266 | 153 |
| | rs34472363 | 0 vs 1 | 843 | 1130 |
| | | 0 vs 2 | 843 | 329 |
| | rs5821892 | 0 vs 1 | 750 | 1150 |
| | | 0 vs 2 | 750 | 414 |
| 20 | rs34472363 | 0 vs 1 | 843 | 1130 |
| | | 0 vs 2 | 843 | 329 |
| 22 | rs2058084 | 0 vs 1 | 891 | 1116 |
| | | 0 vs 2 | 891 | 316 |
| 27 | rs10238953 | 0 vs 1 | 1720 | 565 |
| | | 0 vs 2 | 1720 | 44 |
| | rs2759661 | 0 vs 1 | 1452 | 776 |
| | | 0 vs 2 | 1452 | 101 |
| **Low-freq variants** | | | | |
| 10 | rs151097801 | 0 vs 1/2 | 2325 | 4 |
| | rs117788141 | 0 vs 1/2 | 2325 | 4 |
| | rs140903666 | 0 vs 1/2 | 2319 | 10 |
| | rs144863771 | 0 vs 1/2 | 2319 | 10 |
| | rs147858841 | 0 vs 1/2 | 2324 | 5 |
| 18 | rs142280455 | 0 vs 1/2 | 2324 | 5 |
| | rs137852591 | 0 vs 1/2 | 2324 | 5 |
| 20 | rs143666989 | 0 vs 1/2 | 2324 | 5 |
| | rs148153989 | 0 vs 1/2 | 2325 | 4 |
| 22 | rs149342416 | 0 vs 1/2 | 2323 | 6 |
| | rs143662783 | 0 vs 1/2 | 2325 | 4 |
| 27 | rs142863092 | 0 vs 1/2 | 2325 | 4 |
| | rs137991779 | 0 vs 1/2 | 2324 | 5 |
| 28 | rs142932029 | 0 vs 1/2 | 2326 | 3 |
| | rs201095751 | 0 vs 1/2 | 2325 | 4 |

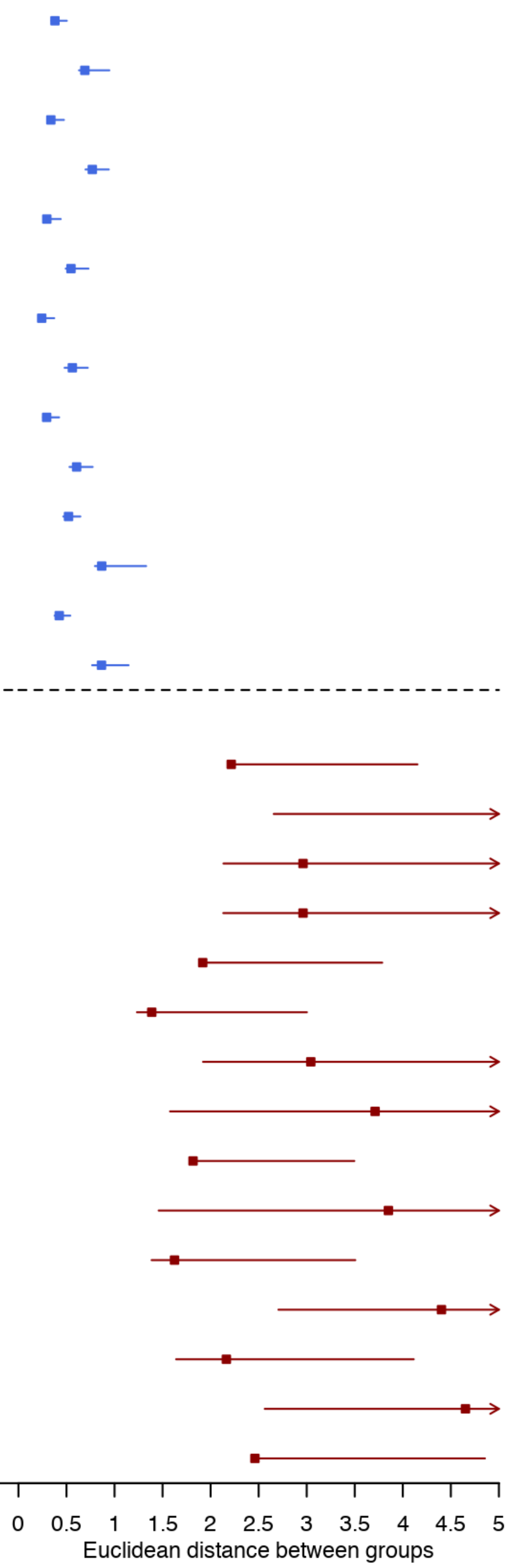


Fig S4. Magnitude of variant effect on facial modules, quantified by the Euclidean distance between averaged faces of different genotype groups. The 95% confidence interval was obtained by 5000 bootstraps. The farther away the blue (common) or red (low-freq) rectangular boxes fall from line x=0, the larger the group distances and the greater the magnitude of effects.  Common variants that yielded significant GWAS association in the same cohort with the same modules are used as a comparison to low-frequency variants. Genotype groups column indicates the two groups of people of whom the faces were averaged and distance was computed. For example, 0 vs 1/2 means minor allele homozygotes vs the remaining. The following two columns indicate sizes of the two groups in comparison. Low-frequency variants had large effects compared to previously reported common variants, although this could be a result from the much smaller size of carrier group and may not reflect genuine greater effects of low-frequency variants.
